# Non-coding Class Switch Recombination-related transcription in human normal and pathological immune responses

**DOI:** 10.1101/384172

**Authors:** Helena Kuri-Magaña, Leonardo Collado-Torres, Andrew E. Jaffe, Humberto Valdovinos-Torres, Marbella Ovilla-Muñoz, Juan M Téllez-Sosa, Laura C Bonifaz Alfonzo, Jesùs Martinez-Barnetche

## Abstract

**Background:** Antibody class switch recombination (CSR) to IgG, IgA or IgE is a hallmark of adaptive immunity, allowing antibody function diversification beyond IgM. CSR involves a deletion of the IgM/IgD constant region genes placing a new acceptor Constant (C_H_) gene, downstream of the VDJ_H_ exon. CSR depends on non-coding (CSRnc) transcription of donor Iμ and acceptor I_H_ exons, located 5’ upstream of each C_H_ coding gene. Although our knowledge of the role of CSRnc transcription has advanced greatly, its extension and importance in healthy and diseased humans is scarce.

**Methods:** We analyzed CSRnc transcription in 70,603 publicly available RNA-seq samples, including GTEx, TCGA and the Sequence Read Archive (SRA) using *recount2*, an online resource consisting of normalized RNA-seq gene and exon counts, as well as coverage *BigWig* files that can be programmatically accessed through R. CSRnc transcription was validated with a qRT-PCR assay for I_μ_, I_γ3_ and I_γ1_ in humans in response to vaccination.

**Results:** We mapped I_H_ transcription for the human IgH locus, including the less understood *IGHD* gene. CSRnc transcription was restricted to B cells and is widely distributed in normal adult tissues, but predominant in blood, spleen, MALT-containing tissues, visceral adipose tissue and some so-called “immune privileged” tissues. However, significant I_γ4_ expression was found even in non-lymphoid fetal tissues. CSRnc expression in cancer tissues mimicked the expression of their normal counterparts, with notable pattern changes in some common cancer subsets. CSRnc transcription in tumors appears to result from tumor infiltration by B cells, since CSRnc transcription was not detected in corresponding tumor-derived immortal cell lines. Additionally, significantly increased I5 transcription in ileal mucosa in Crohn’s disease with ulceration was found.

**Conclusions:** CSRnc transcription occurs in multiple anatomical locations beyond classical secondary lymphoid organs, representing a potentially useful marker of effector B cell responses in normal and pathological immune responses. The pattern of IH exon expression may reveal clues of the local immune response (i.e. cytokine milieu) in health and disease. This is a great example of how the public *recount2* data can be used to further our understanding of transcription, including regions outside the known transcriptome.

## Background

The hallmark of the humoral adaptive immune response is the production of high affinity class-switched antibodies from relatively low-affinity IgM^+^ naive precursors. Affinity maturation and class switch recombination (CSR) are tightly regulated, molecular processes that occur in a specialized microenvironment within secondary lymphoid organs known as the germinal center (GC). Upon T-dependent antigen stimulation, IgM^+^ naive B cells re-localize into the B cell follicle, undergo clonal expansion within the GC. The GC reaction is an iterative process of mutation-selection that leads to the generation of antigen-specific high affinity, class-switched memory B cells and long-lived antibody secreting plasma cells (reviewed in [1]). Antibody class switching from IgM to IgG, IgA or IgE allows effector function diversification. Both memory B cells and long-lived plasma cells are critical determinants of vaccine efficacy [2].

CSR involves a deletion of a genomic segment from the *IGHM/IGHD* coding interval (C_μ_-C_δ_) to the upstream flank of one of the *IGHG, IGHA* or *IGHE* genes in the telomeric region of human chromosome 14. Activated GC B cells upregulate the Activation Induced Cytidine Deaminase (AID), which deaminates cytidines in the G-rich Switch (S) regions located upstream of each immunoglobulin constant coding gene (C_H_). Cytidine deamination induces DNA damage response, which eventually leads to double stranded DNA breaks in both donor (S_μ_) and the corresponding acceptor S region. The chromosomal ends are rejoined and the C_μ_-C_δ_-encoding intervening DNA segment is re-circularized in a non-replicating episome by non-homologous end joining (reviewed in [3]).

The initiation of CSR depends on non-coding transcription of I_H_ exons, known as germline or “sterile” transcripts (referred hereafter as CSRnc transcription). I_H_ exons are located in the 5’ region of each S-C_H_ gene module. Non-coding transcription of I_H_ exons extends to the S and CH region, is coupled to chromatin remodeling and is dependent on splicing [4, 5]. CSRnc transcripts form an R-loop in the corresponding S region, which recruits AID to target S region deamination and CSR (reviewed in [6]). The precise mechanism of AID targeting to the S_H_ region remains elusive, and off-target AID activity is implicated in the genesis of B cell malignancies [6, 7].

CSR is a complex cellular process that occurs in specialized microenvironments in secondary and tertiary lymphoid organs. The cellular choice of which I_H_ to transcribe, and consequently the Ig class to switch to, is influenced by the availability of certain cytokines such as IL-4, IFNγ, TGFβ and P AMP’s, among others. Such environmental cues are thought to trigger specific signals that promote selective transcription of a given I_H_ exon, guiding CRS according to a particular microenvironment or pathogenic insult [3]. CSRnc transcription patterns may reflect distinctive immunological events, such as the dependence of T cell help and other micro-environmental signals. Thus, CSRnc transcription quantitation during normal and pathological human immune responses could uncover novel pathogenic mechanisms and transcriptional signatures with potential clinical value. In addition, despite CSRnc transcription is biologically linked to B cells, its expression in other cell types has not been ruled out.

The recent explosion in the generation of public genomic data, and in particular transcriptome-wide profiling with RNA sequencing (RNA-seq) provides a unique opportunity to explore previously unannotated features in the human genome. To characterize CSRnc transcription in normal and pathological conditions, we tested CSRnc transcription in human vaccination and analyzed the transcriptional landscape of the human IgH locus using more than 70,000 publicly available human RNA-seq samples from a wide variety of research projects, including the Genotype Tissue Expression project (GTEx) [8, 9], The Cancer Genome Atlas (TCGA) [10, 11], and more than 2,000 projects from the Sequence Read Archive (SRA) using *recount2* [12].

## Results

### CSRnc transcription is B cell-specific and its boundaries are diffuse

Overall, *recount2* [12] comprises a highly heterogeneous catalog of RNA-seq experiments belonging to 2,036 independent studies and comprising 70,603 samples. Each study is composed of an average of 34 samples. However, TCGA [10, 11] and GTEx (SRP012682) [8, 9] are two projects (studies) with the largest number of sequencing samples (11,284 and 9,661 respectively), and represent 29 % of our dataset (**Additional file 1: Figure S1**).

Although human CSRnc transcription has been evaluated by RT-PCR [5, 13, 14], the precise transcription boundaries, including alternate transcription initiation sites and splicing variants remain undefined. We selected RNA-seq samples derived from normal FACSorted B cells to map CSRnc transcription and to define transcriptional boundaries for read count quantitation for further analysis (**Additional file 2: Table S1**) [15-21]. We found that in normal adult B cells, CSRnc boundaries are less sharply defined than coding transcripts, and as expected, extend into switch regions (Figure 1) [5]. Projects SRP045500 [18] and SRP051688 [20] describing the transcriptome of isolated peripheral blood (PB) immune cells in healthy adults, including primary neutrophils, monocytes, myeloid dendritic cells, B, NK and T cells revealed that among terminally differentiated hematopoietic lineage-derived cells, CSRnc transcription is restricted to B cells (Figure 1A, **Additional file 1: Figure S2**).

**Figure 1.**
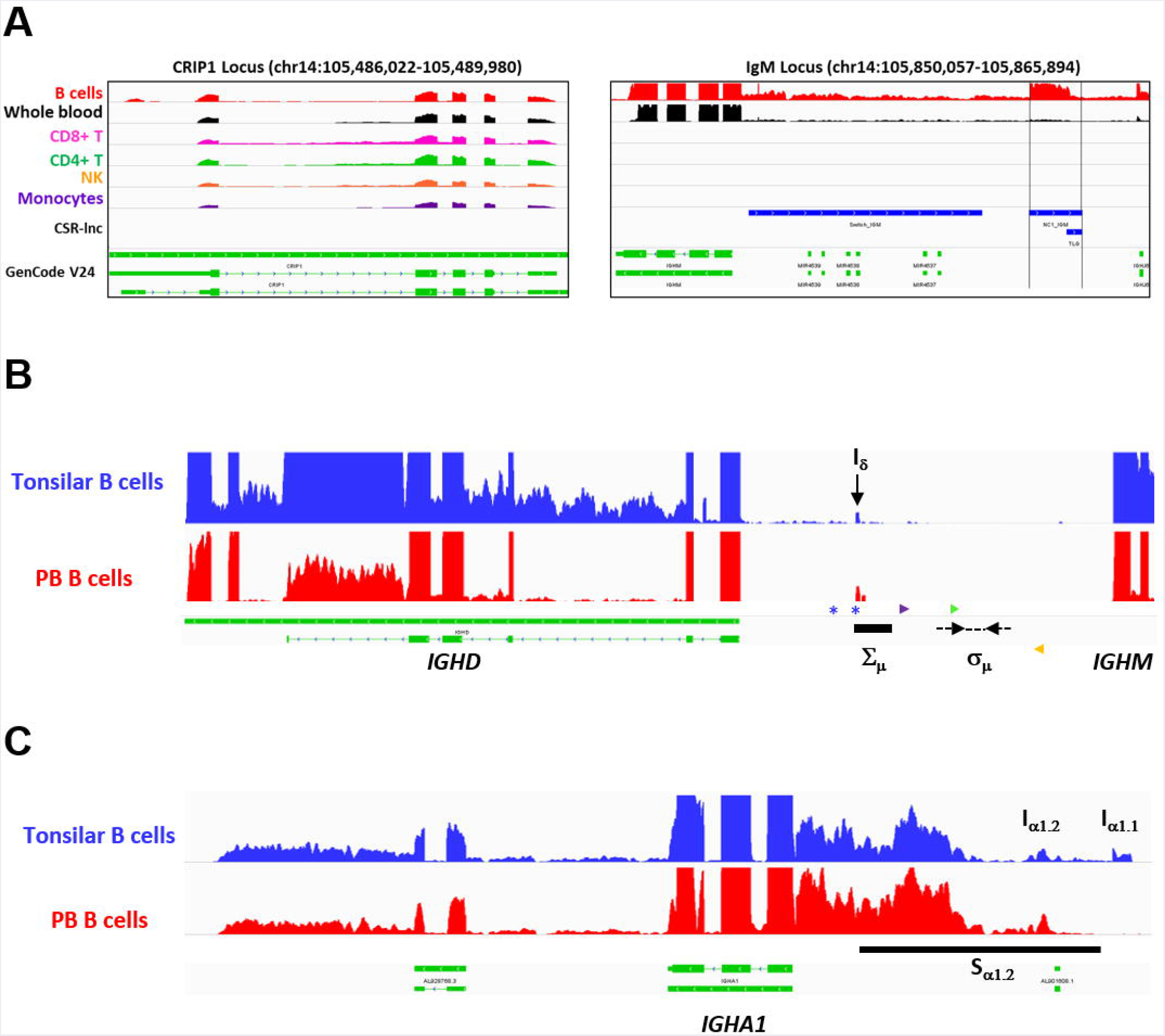
CSRnc transcription boundaries definition. Selected projects using isolated B cells were used to define the limits of CSRnc transcription in human chromosome 14 (See **Additional file 2: Table S1**). The telomeric region is towards the right. *BigWig* files were downloaded using *recount2* and visualized using IGV [75] to inspect to determine the boundaries for each CSRnc transcript. **A**) Coverage graph of project SRP045500 showing the CRIP1 locus *(left panel)* and the *IGHM* locus *(right panel)* in B *(red track)* and non-B cells. The CRIP1 gene is transcribed in B and non-B cells, whereas *IGHM* and I_μ_ (within black vertical lines) is transcribed only in B cells and in peripheral blood *(Black track*). Predicted S_μ_ region is shown in the bottom track *(blue)* and annotated genes (GenCode V24) are shown in *green*. **B**) A view of the *IGHM – IGHD* intron locus displaying coverage graphs from normal purified naive tonsillar (SRP021509, blue track) and peripheral blood CD19+ B cells (SRP060715, red track). The vertical black arrow shows I_δ_. *IGHD* annotation GenCode V24 is shown in *green*. The I_δ_ exon overlaps with the Σμ region [22, 43, 45] and is centromeric regarding the mapped sites for μ – δ CSR junctions (Dotted black arrows) [43]. Antisense primer (18156) used by Kluin *et al*. [43] and Arpin, *et al*. (P4) [45] are shown in purple and green, respectively. The S_δ_ Sense primer used by Chen, *et al*. [44] is shown in orange. The *blue asterisks* indicate *Hind* III sites [22, 23, 43]. **C**) Transcriptional landscape of the *IGHA1* locus displaying coverage graphs of the *IGHA1*, GenCode V24 annotation *(green track)*, tonsilar naive (SRR834982, blue) and peripheral blood B cells (SRR2097501, red). Both I_α1.1_ and I_α1.2_ transcripts are shown. The S_α1_ region *(black bar)* is shown.

CSRnc and C_H_ coding transcription in isolated PB CD19C B cells was analyzed. High relative transcription of Iμ and Cμ, intermediate relative transcription of I_μ_, I_γ3_, I_γ1_, I_α1_._2_, I^γ2^, I_γ4_ and I_α2_, and low transcription of I_ε_ and I_α1.1_ were characteristic of PB B cells (**Additional file 1: Figure S3A-C**) [15, 18]. Furthermore, CSRnc transcription was analyzed in tonsillar naive and germinal center B cells from project SRP021509 [16]. CSRnc transcription for most I_H_ but not I*ε*, was relatively high in both naive and GC B cells (**Additional file 1. Figure S3D**). These findings indicate that CSRnc transcription is not exclusive of activated B cells, and agrees with previous findings demonstrating constitutive CSRnc transcription [5, 14].

A transcriptionally active 309 base-pair (bp) region within the *IGHM-IGHD* intron was identified (referred hereafter as I_δ_) and was included for further analysis (Figure 1B). This region is homologous to Iμ, overlaps with a previously described repeat termed Σμ implicated in μ-δ CSR in IgD^+^ myelomas [22, 23]. For the *IGHA1* gene, we identified two potential I_H_ exons (I_α1.1_ and I_α1_._2_) that were selected for downstream analysis (Figure 1C). I_γ2_ overlapped with a previously annotated lincRNA, ENST00000497397.1 (AL928742.1) at ENSEMBL [24]. The genome coordinates of each I_H_ exon identified and further analyzed are in Table 1. Navigation across the IgH locus using the ENSEMBL Genome Browser [24] allowed to confirm that the predicted I_H_ exons include regions of RNApolII, H3K36 and H3K4 trimethylation enrichment in peripheral blood B cells and EBV-transformed B cells generated by the Roadmap Epigenomics and ENCODE projects [25, 26], indicating active transcription (**Additional file 1: Figure S4**). Overall, these results agree with the current model of CSRnc transcription [5]; however, the identification of novel transcribed elements ads complexity to the transcriptional regulation of the IgH locus.

**Table 1.** Mapping I exons (IH) and switch regions (S) in human
chromosome 14.GRCh38.p10

|  |  |  |  |
| --- | --- | --- | --- |
| **Gene** | **Start** | **End** | **Length (bp)** |
| ***IGHM*** |  |  |  |
| Iμ | 105,861,311 | 105,862,213 | 903 |
| Sμ | 105,856,501 | 105,860,500 | 4,000 |
| ***IGHD*** |  |  |  |
| Iδ | 105,847,549 | 105,847,857 | 309 |
| ***IGHG3*** |  |  |  |
| I$\gamma_3$ | 105,775,023 | 105,775,702 | 680 |
| S$\gamma_3$ | 105,773,001 | 105,774,000 | 1,000 |
| ***IGHG1*** |  |  |  |
| I$\gamma_1$ | 105,747,322 | 105,748,299 | 978 |
| S$\gamma_1$ | 105,744,501 | 105,745,500 | 1,000 |
| ***IGHA1*** |  |  |  |
| I$\alpha_{1.2}$ | 105,711,304 | 105,711,907 | 604 |
| I$\alpha_{1.1}$ | 105,712,639 | 105,712,892 | 254 |
| S$\alpha_1$ | 105,709,501 | 105,712,500 | 3,000 |
| ***IGHG2*** |  |  |  |
| I$\gamma_2$ | 105,647,803 | 105,648,549 | 747 |
| S$\gamma_2$ | 105,627,501 | 105,628,500 | 1,000 |
| ***IGHG4*** |  |  |  |
| I$\gamma_4$ | 105,629,000 | 105,629,500 | 501 |
| S$\gamma_4$ | 105,623,001 | 105,624,000 | 1,000 |
| ***IGH E*** |  |  |  |
| Iε | 105,605,036 | 105,605,357 | 322 |
| Sε | 105,602,501 | 105,604,500 | 2,000 |
| ***IGHA2*** |  |  |  |
| I$\alpha_2$ | 105,589,300 | 105,590,300 | 1,001 |
| S$\alpha_2$ | 105,589,001 | 105,591,500 | 2,500 |

### CSRnc transcription in peripheral blood is modified upon vaccination and does not depend on circulating plasmablasts

CSRnc transcription increases upon B cell activation [5]. To validate that the predicted CSRnc transcripts were indeed induced upon activation, normal human B cells were stimulated with agents mimicking T-dependent activation (CD40 ligand, IL-21 and CpG) and T independent activation (CpG, Pokeweed mitogen and SAC) *in vitro* for 3 and 6 days. Total RNA was obtained and I_μ_, I_γ3_ and I_γ1_ CSRnc and AID transcripts were quantified by qRT-PCR (Figure 2A-D). CSRnc transcripts were detected in B cells activated by both T-dependent and T-independent activators, but transcription levels were significantly higher for T-dependent like activation. In both types of activation, CSRnc transcription at 6 days post-activation was higher than 3 days post-activation (Figure 2A-D). The highest transcription was for Iμ (3-fold higher than I_γ1_ and I_γ3_). Transcription of AID showed the same pattern as CSRnc transcripts (Figure 2D).

**Figure 2.**
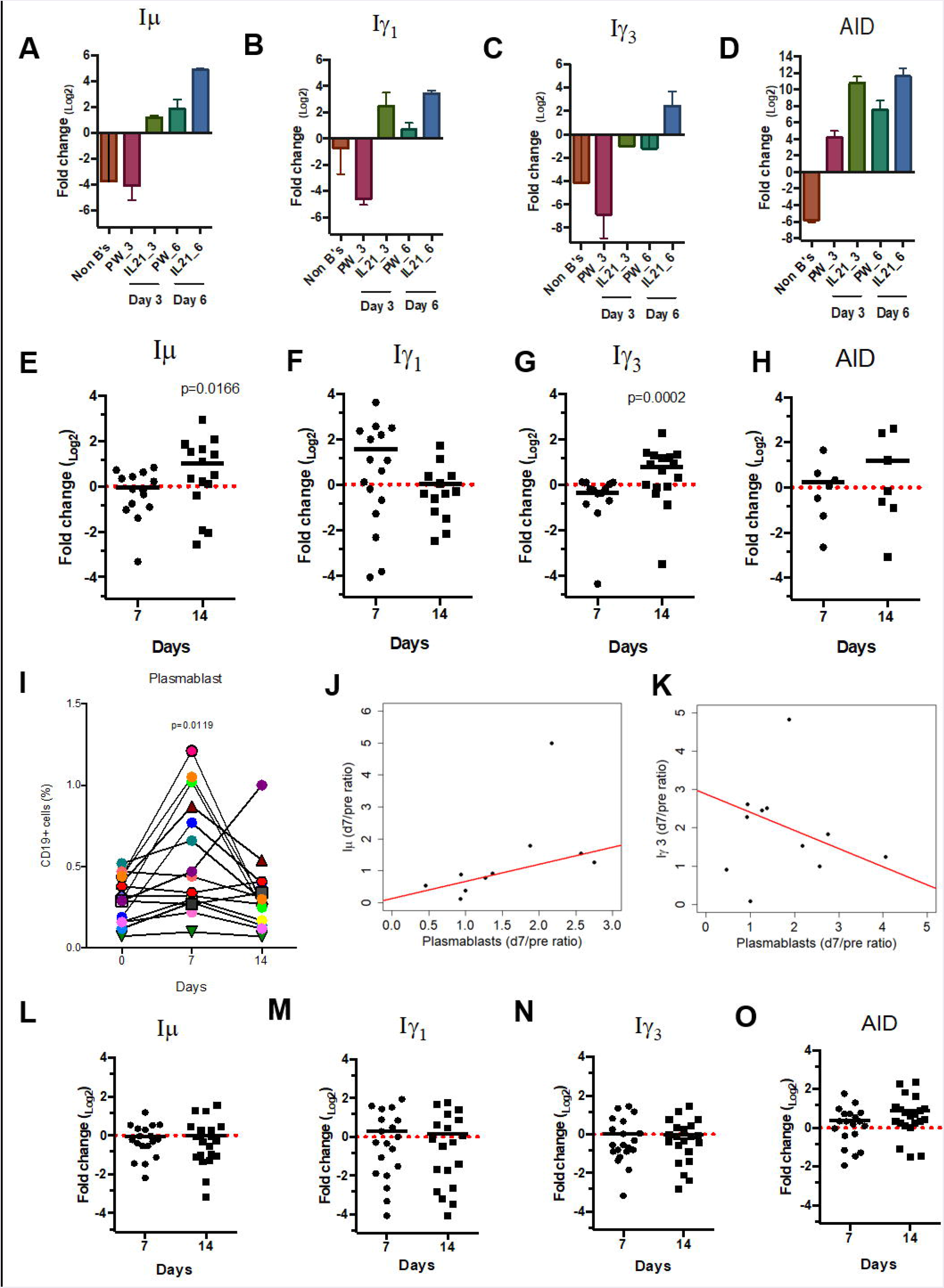
CSRnc transcripts quantitation *in vitro* and in vaccination. A qPCR Taq-Man assay for I_μ_, I_γ3_ I_γ1_ and AID was used to quantitate CSRnc transcription with the 2^ΔΔCt^ method (log2). **A-D**) Enriched B cells cultured for 3 and 6 days with T-dependent like activation (IL-21, a-CD40 and CpG) and T-independent activation (PWM, SAC and CpG). Bar plots represent the mean fold-change (non-activated enriched B cells /activated B cells) of two independent experiments. Non-B cells were used as control. **E-H**) qPCR Taq-Man assay for I_μ_, I_γ3_ I_γ1_ and AID from total RNA obtained from donors’ PBMCs taken at preimmunization (day 0) against Hepatitis B and/or Tetanus-Diphteria and on days 7 and 14 post-immunization. (Wilcoxon test. p<0.05). **I**) Plasmablast (CD3’CD19+CD20’ CD27+CD38+) mobilization in peripheral blood expressed as a percentage of CD 19_+_ B cells (Wilcoxon test. p =0.005). J) Positive correlation between day 0/7 plasmablast ratio and day 0/7 I_μ_ ratio (LTS regression method. Adjusted R^2^: 0.53, p-value: 0.015). K) Negative correlation between day 0/7 plasmablast ratio and day 0/14 I_γ3_ ratio (LTS regression method. Adjusted R^2^ = 0.63, p-value: 0.011). **L-O**) No significant changes in CSRnc transcription assessed by qPCR were observed 7 and 14 days after trivalent inactivated Influenza vaccination (Wilcoxon test. P > 0.05). **L**) I_μ_, **M**) I_γ3_, **N**) I_γ1_ and **O**) AID. Dotted red line indicates no change in expression (Fold-change = 1.0).

Immunization promotes an antigen-specific mobilization of plasmablasts to peripheral blood around day 7 post-challenge [27, 28]. This plasmablast wave is thought to derive from germinal centers [29]. Thus, we hypothesized that the level of CSRnc transcription in peripheral blood correlates with the amount of plasmablasts. We assessed CSRnc transcription and plasmablast proportion in peripheral blood of healthy subjects before, 7 and 14 days post vaccination with either hepatitis B (HB) alone or in combination with tetanus/diphtheria vaccine (TT/Dp) (Figure 2E-K). Regarding pre-immune levels, Iμ transcription was not affected at day 7 (mean 0.95 ± 0.5, p = 0.76) or at day 14 (mean 2.04 ± 2, p = 0.073), however at day 14 was higher than at day 7 (p = 0.01) (Figure 2E). I_γl_ transcription was neither affected at day 7 (mean 2.9 ± 3.2, p = 0.057) nor at day 14 (mean 1.02 ± 0.9. p = 0.57) (Figure 2F). Contrastingly, I_γ3_ transcription regarding pre-immune levels was reduced at day 7 (mean 0.76 ± 0.3, p = 0.016) and increased at day 14 (mean 1. 73 ± 1.1, p = 0.021). As expected, I_γ3_ transcription at day 14 was higher than at day 7 (p =0. 0002) (Figure 2G). No changes were detected in AID expression (Figure 2H).

Plasmablast levels peaked at day 7 post-vaccination (p = 0.011), and returned to pre-vaccine levels 14 days post-vaccination, as previously described [27, 28]. There was no correlation between plasmablasts and I_γ1_ transcription increase at day 7 (LTS method. Adjusted R^2^ = -0.1265, F-statistic: 0.1013 on 1 and 7 DF, p = 0.75). However, plasmablast and Iμ transcription correlated at day 7 (Figure 2J) (LTS method. Adjusted R^2^ = 0.53; F-statistic = 10.17 on 1 and 7 DF, p = 0.015). Interestingly, plasmablast fold-change at day 7 negatively correlated with I_γ3_ transcription (Figure 2K) (LTS method. Adjusted R = 0.63; F-statistic = 13.04 on 1 and 6 DF, p-value: 0.011). No changes in I_μ_, I_γ1_ and I_γ3_ transcription was detected in response to influenza vaccination (Figure 2L-O). Overall, these results suggest that an increase in CSRnc transcription in peripheral blood upon vaccination is dependent on vaccine type, and that the contribution of vaccine-mobilized plasmablasts to CSRnc transcription is negligible.

### CSRnc transcription is prevalent in a large fraction of the *recount2* dataset, with predominant transcription of I_μ_

Normalized counts (RPKM) obtained with *recount2* were used to asses CSRnc transcription in the SRA, TCGA and GTEx datasets. We found no CSRnc transcription (RPKM = 0) in any of the 10 I_H_ exons in a substantial fraction of the whole dataset (n = 26,512 samples; 37%) (Table 2). The I_H_ log2-transformed RPKM average (2.65) for each non-zero RPKM sample (n = 44,091; 62.4 %) was used to define an expression cutoff. “High” CSRnc transcription was defined as a mean log_2_ RPKM of 2.65 or higher. Only 29.8 % of RNA-seq samples (n = 21,017) were above this expression cutoff. “Low” CSRnc transcription was defined as a mean log_2_ RPKM < 2.65 (Figure 3A. Table 2), which accounted for the remaining 32.7 % of the samples (n = 23,074). The CSRnc transcription levels varied according to I_H_. Higher transcription (log_2_ RPKM), as well as a more widespread transcription (proportion of samples) was found for Iμ in all datasets (**Additional file 1: Figure S5**).

**Table 2.** General sequencing run metrics used for CSR-ncRNA expression analysis

|  | **log<sub>2</sub>RPKM** | **Samples** | **(%)** |
| --- | --- | --- | --- |
| Total | NA | 70,603 | **100.0** |
| Highly expressed | > 2.65 | 21,017 | **29.8** |
| Low-expressed | < 2.6, > 0 | 23,074 | **32.1** |
| Not expressed | 0 | 26,512 | **37.1** |
| Clustering set | > 0 | 44,091 | **62.9** |

**Figure 3.**
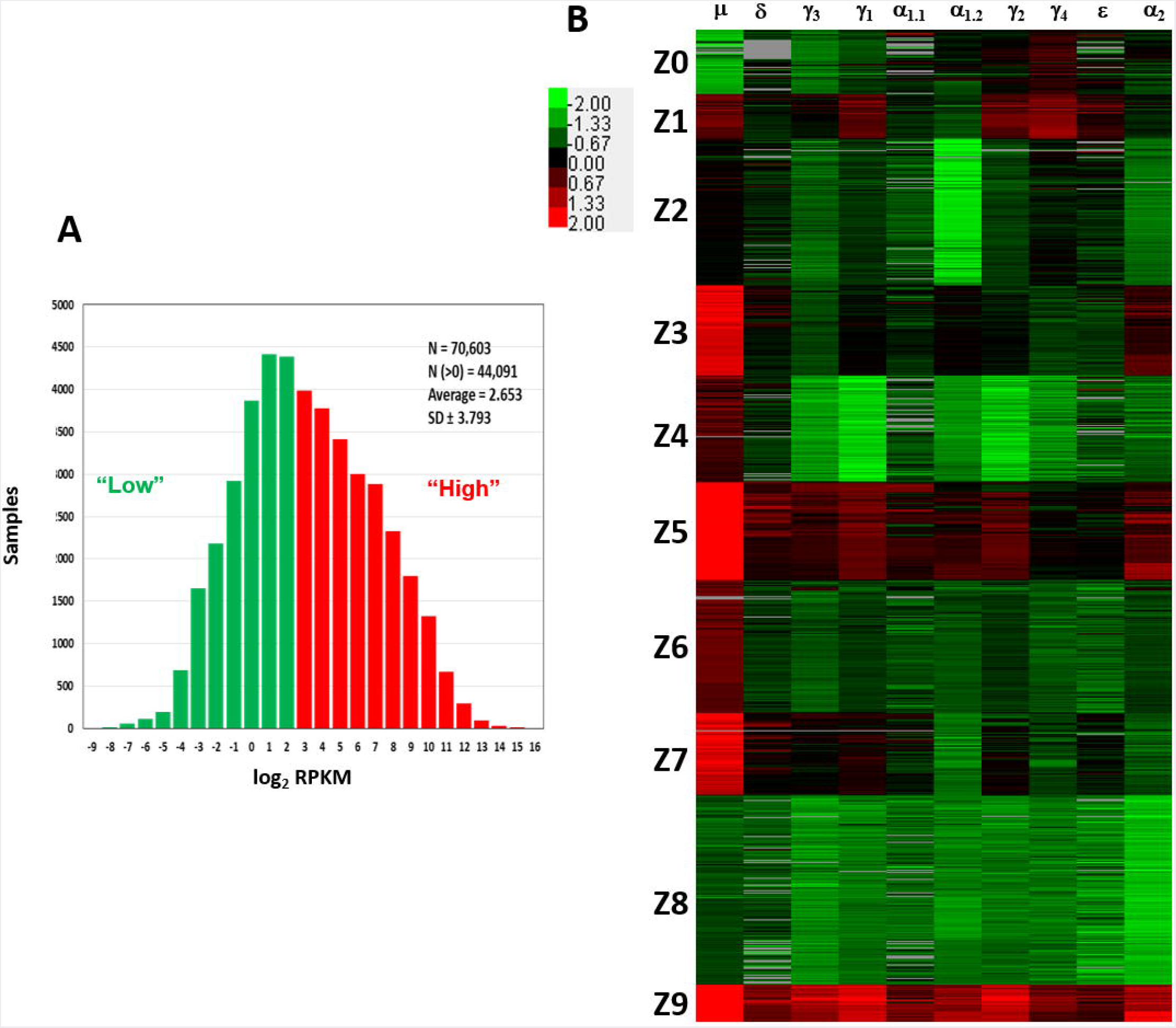
Quantitative analysis of CSRnc transcription using *recount2*. **A**) Average log_2_-transformed RPKM distribution of the 10 I_H_ exons per sample. There were 44,091 samples with non-zero RPKM. Higher than the mean (log_2_RPKM > 2.65) was considered “highly” transcribed (shown in red). “Low” CSRnc transcription was defined as a mean log_2_ RPKM < 2.65 (green). **B**) CSRnc transcription profiling by *Z*-score clustering. Log2RPKM CSRnc transcription values of GTEx + TCGA + SRA datasets were transformed to standardized Z-scores and subjected to *k*-means clustering with a predefined number of 10 clusters (*Left panel*). Clustered data is represented in a heatmap where I_H_’s are columns and Z clusters are in rows. Negative Z-scores (i.e. < 2.65 log_2_RPKM) are shown in green, positive Z-scores (i.e. > 2.65 log2RPKM) are shown in red. Absent values (RPKM = 0) are shown in grey. Z values near 0 are shown in black. The pattern expression of each cluster is represented in boxplots in **Additional file 1: Figure S6**.

#### The *recount2* dataset is partitioned in distinctive CSRnc transcription profiles

We further de-convoluted CSRnc transcription according to I_H_ relative transcription profile. To do so, the entire non-zero dataset including GTEx, TCGA and SRA Z-scores was clustered in 10 groups using *k*-means clustering (**Figure 3B. Additional file 1: Figure S6**). Using Z0 as reference, each of I_H_ showed a distinctive expression pattern in the remaining Z clusters (log_2_ RPKM regression analysis. F-statistic: p-value: < 2.2e-16). Clusters Z0, Z2 and Z8 were characterized by low (mean log_2_ RPKM < 2.65 or Z score < 0) CSRnc transcription in all I_H_ classes. Clusters Z4, Z6 and Z7 were characterized by “high” expression of Iμ only (mean log_2_RPKM > 2.65). Cluster Z3 showed “high” expression in I_μ_ and I_α2_. Cluster Z1 showed high expression in I_μ_, I_γ1_ and I_γ4_. Finally, clusters Z5 and Z9 showed “high” expression in all I_H_’s (**Figure 3B. Additional file 1: Figure S6**).

### CSRnc transcription is widely distributed in healthy tissues, with particular profiles according to tissue

To gain insight into CSRnc expression patterns in healthy adult human tissues, we used GTEx samples as a reference. Non-zero RPKM per tissue samples were ranked according to their mean log_2_ RPKM. Higher average transcription was found in lymphoid tissues such as spleen, EBV-transformed B lymphocytes and whole blood, but also in organs with mucosal-associated lymphoid tissues (MALT) such as terminal ileum, transverse colon, stomach, lung and esophageal mucosa (Figure 4). A remarkable difference in average log_2_ RPKM is observed in transverse colon (mean 7.5 ± 3.9) and sigmoid colon (mean 2.1 ± 3.2). Interestingly, salivary gland expression was among the tissues with highest transcription, and non-mucosal tissues such as thyroid and pituitary gland showed high average I_H_ transcription. Another notable difference in average log_2_ RPKM was observed between visceral adipose tissue (omentum; mean 4.9 ± 2.3) and subcutaneous adipose tissue (mean 1.9 ± 1.5). Conversely, samples of tissues such as brain, skeletal and cardiac muscle, skin, as well as chronic myelogenous leukemia cell line K562 and transformed dermal fibroblast cell lines showed the lowest CSRnc transcription levels (mean log_2_ RPKM < 2.65) (Figure 4).

**Figure 4.**
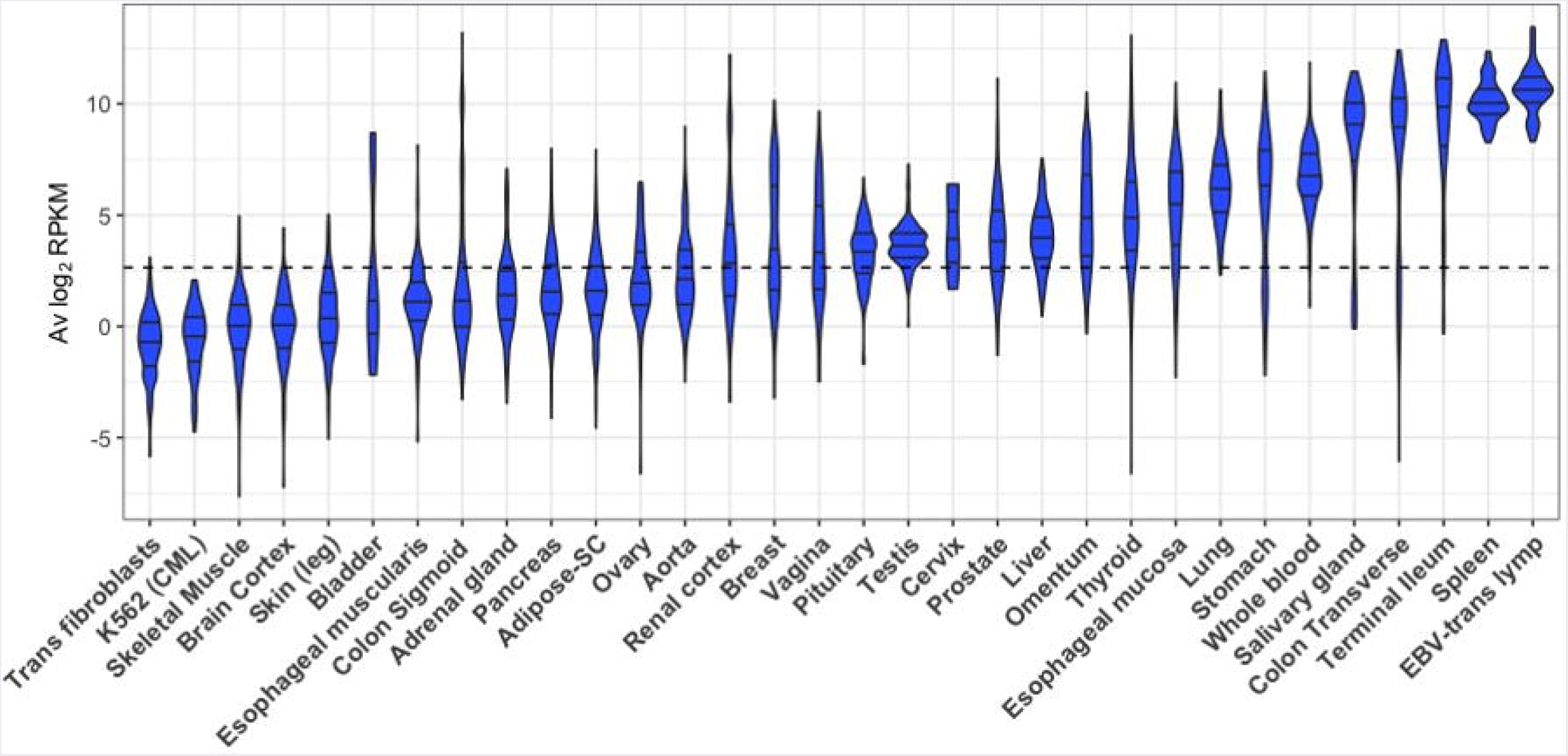
CSRnc transcription in healthy adult tissues. The GTEx dataset was partitioned according to tissue type. Violin plot of average log_2_RPKM distribution of GTEx project representative tissues, ordered form left to right according to each tissue median log_2_RPKM. The violin area was scaled according to sample count and median and quartiles are shown. A dotted black line marks the mean average log_2_RPKM (2.65). For simplicity, only brain cortex was included as a representative sample for central nervous system.

Using whole blood as reference tissue, we performed a linear regression analysis of each I_H_ by tissue type. The transcription pattern of each I_H_ in whole blood was significantly different (p < 2.2e-16) (Figure 5). Iμ transcription was similar to I_H_ average transcription (Figure 4 and 5), in which higher transcription was observed in spleen, terminal ileum, salivary gland, and transverse colon than blood. Of note, Iμ transcription in testis was particularly low, despite its high average transcription. I_δ_ transcription was similar to I_μ_ transcription, but only spleen and terminal ileum were significantly higher. I_γ_, I_γ1_ and I_γ2_ transcription was similar and was higher in spleen than in blood (p < 2e-16) (Figure 5).

**Figure 5.**
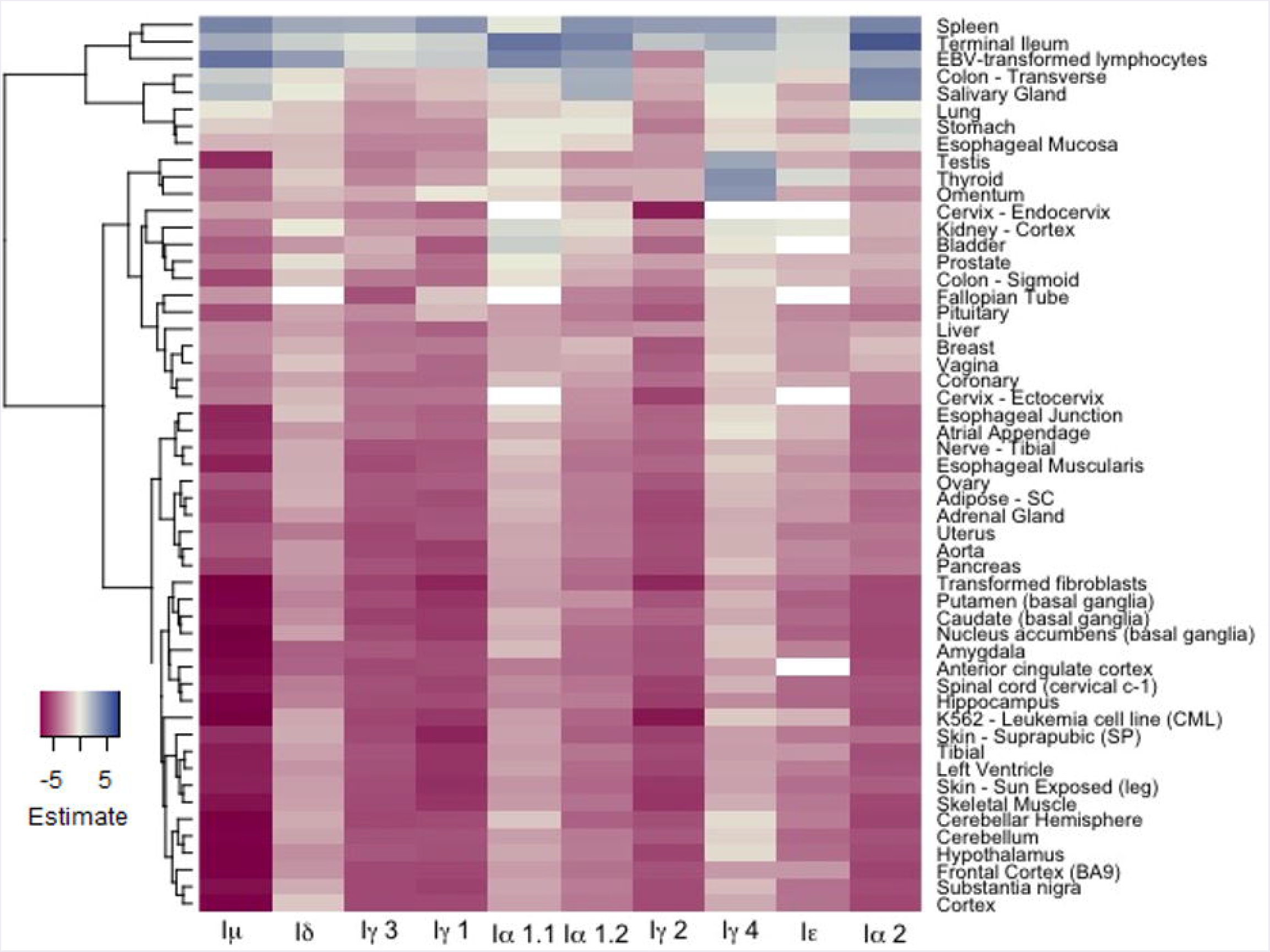
Regression analysis of CSRnc transcription as a function of tissue type and I-exon. Whole blood CSRnc transcription was used as reference tissue (*rows*) for regression analysis according to I-exon (*columns*). The estimate for each comparison is expressed as a heatmap. Higher CSRnc transcription than in blood is represented in blue tones, whereas lower transcription is shown in pink tones. Zero estimate values are shown in ivory. Missing values (NA’s) are shown in white. Euclidian distance was used for hierarchical clustering by row.

For most tissues, transcription of I_α1.1_ and I_α1.2_ was highly correlated, however some differences were noted. I_α1.1_ transcription was higher in terminal ileum and transverse colon than in blood (p < 0.001). In contrast, I_α1.2_ transcription was higher in spleen and salivary gland than in blood (p < 2e-16). I_α2_ transcription was similar to I_α1.2_ transcription, but also was higher in stomach, esophageal mucosa than in blood (p < 1.1e-05) (Figure 5). Thus, I_ρ1_ and I_ρ2_ transcription pattern matches with the fact of IgA as the main immunoglobulin in mucosal tissue. Furthermore, our results suggest that CSR to IgA may involve tissue-dependent alternative transcription initiation and/or splicing in the corresponding CSRnc transcripts.

The most unexpected patterns of CSRnc transcription corresponded to I_γ4_ and I_ε_. Transcription for both was higher in spleen, terminal ileum and EBV-transformed B-cells than in blood (p < 0.01). However, I_γ4_ transcription was higher in thyroid, visceral adipose tissue (omentum), testis, than in blood (p < 2e-16). Similarly, I_ε_ transcription was higher in in thyroid than in blood (p < 0.00003) (Figure 5).

Consistently, de-convolution of CSRnc transcription according to I_H_ relative expression (Z-score) revealed that terminal ileum, spleen, transverse colon, whole blood and salivary gland share a similar expression pattern and are highly enriched (FDR < 0.001) in clusters Z5 and Z9 (high transcription in all I_H_’s) (Figure 6). Some tissues with MALT such as stomach, esophageal mucosa and lung shared a similar I_H_ transcription pattern and were enriched in cluster Z5 and Z3 (high I_μ_ and I_α2_ transcription) (FDR < 0.001), whereas others such as breast, vagina, liver were enriched in cluster Z3 and Z6 (I_μ_ only). Testis, thyroid, pituitary and omentum, characterized by the highest I_γ4_ transcription, were enriched in cluster Z1 (I_μ_, I_γ1_ and high I_γ4_) (FDR < 0.001). Finally, the remaining tissues such as subcutaneous adipose tissue, arteries, brain, skeletal and cardiac muscle, skin, transformed fibroblasts and K562 cell lines, were enriched in Z0, Z2 and Z8 (low CSRnc transcription in all I_H_ classes) (FDR < 0.001) (Figure 6), and in few cases were enriched in clusters Z4 and Z6 (I_μ_ only) (FDR < 0.001).

**Figure 6.**
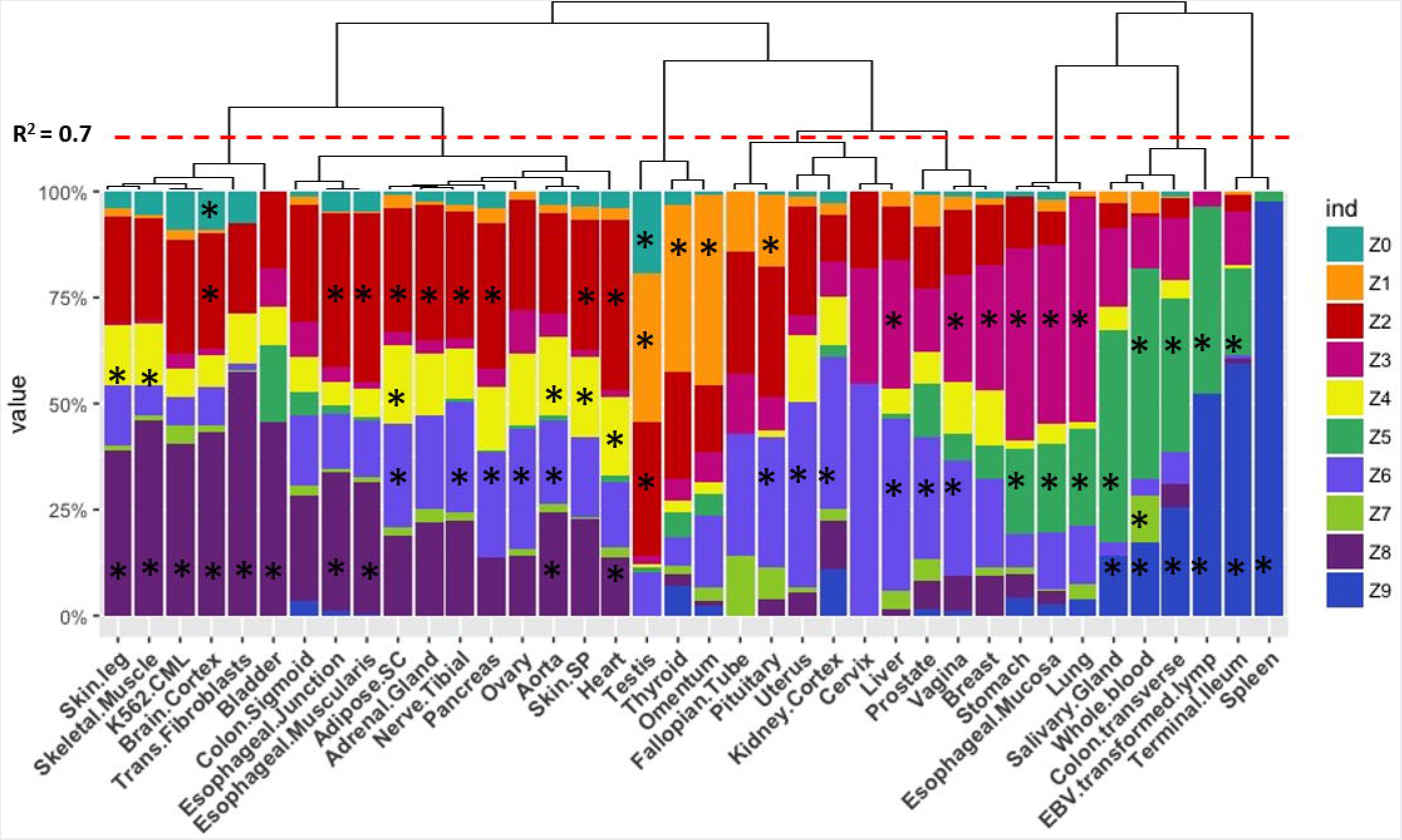
CSRnc transcription profile variation in healthy tissue. The GTEX dataset was categorized according to tissue (*x* axis) and the relative proportion of samples (y-axis) belonging to each Z cluster (in *colors*). Many central nervous system tissues with highly similar pattern were removed for simplicity. Proportions were hierarchically clustered using Pearson correlation as distance metric with Cluster3.0 [75]. The red dotted line marks a correlation R^2^ = 0.7. The black asterisk indicates a significant enrichment of a given tissue in a particular Z cluster (Exact Fisher’s test, Benjamini-Hochberg adjustment for multiple correction. FDR < 0.01).

The observed anatomical distribution and abundance of CSRnc transcription suggests that the amount of CSRnc transcription may be dependent on the abundance of B cells present in a given tissue. We used the C_H_ coding transcript log_2_RPKM as a proxy of the amount of B cells in the tissue. In general, C_H_ transcription was 10 to 100-fold higher than CSRnc transcription. To correlate CSRnc transcription with corresponding C_H_ transcription, Z-scores were used. As expected for each class, CSRnc and C_H_ Z-scores where significantly correlated (p < 1.0e-16), suggesting that the higher numbers of B cells or plasma cells in a given tissue, higher CSRnc transcription. However, we have noticed that for most classes, a fraction of samples deviates from the expected orthogonal correlation, indicating higher CSRnc transcription relative to the C_H_ transcript. This is particularly notable for I_μ_, but also, I_γ1_, I_γ4_ and I_α2_ (**Additional file 1: Figure S7**).

### Iμ. transcription occurs in early lymphoid progenitors and I_γ4_ is widely expressed in non-lymphoid fetal tissues

CSRnc transcription was addressed in early lymphoid development using data from study SRP058719, which addresses transcription in early lymphoid differentiation prior to and after B and T cell lineage commitment using RNA-seq from FACSorted cells [30]. Interestingly, both B and T lineage precursors expressed I_μ_. Enriched hematopoietic stem cells (HSC’s CD34^+^CD38^−^lin^−^), lymphoid-primed multipotent progenitors (LMPP’s, CD34+CD38+CD10^−^CD45RA^+^lin^−^), common lymphoid progenitors (CLP’s, (CD34^+^CD38^+^CD10^+^CD45RA^+^lin^−^), thymic CD34^+^CD7 CD1a CD4 CD8 (Thy1) precursors and fully B cell-committed progenitors (BCPs, CD34^+^CD38^+^CD19^+^) expressed I_μ_ (**Additional file 1: Figures S8 and S9**).

CSRnc expression was analyzed in fetal tissue by using project SRP055513 data, which reported an extensive RNA-seq analysis in twenty fetal tissues during gestational weeks 9-22 [31]. Remarkably, a robust I_γ4_ expression was detected in all fetal tissues tested, regardless of the gestational week and in the absence of additional I_H_ and coding CH transcription (Figure 7). Higher average I_γ4_ expression is in spleen, followed by lung and liver. The latter has lympho-hematopoietic function in the fetal stage. In contrast with their adult counterparts, I_γ4_ was highly transcribed (Z-score > 0.8) in kidney, brain and muscle.

**Figure 7.**
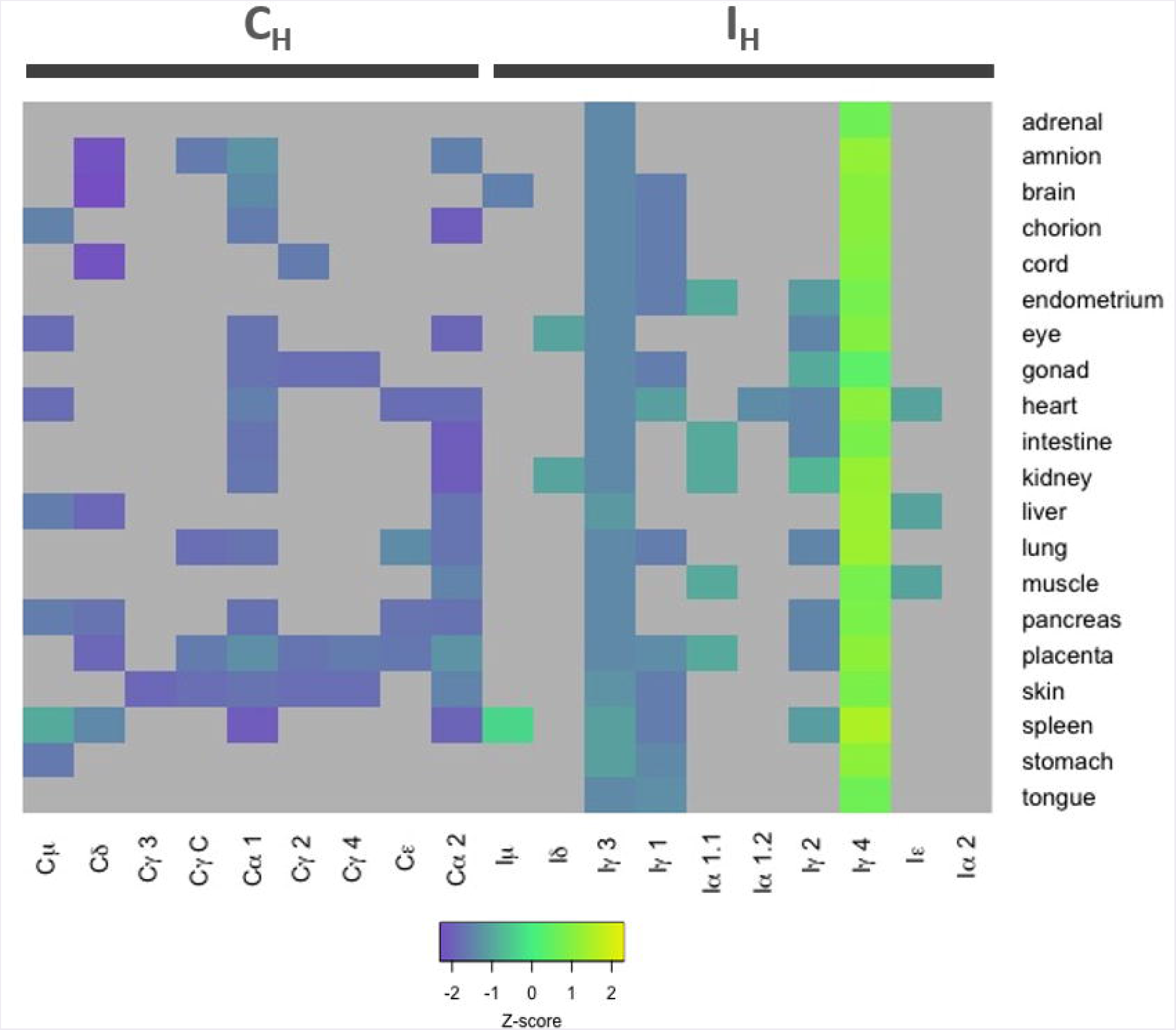
CSRnc I_H_ and Ch transcription in fetal tissues. Heatmap of Z-score average per tissue (n = 3-8) of I_H_ and C_H_ *(columns)* in 9-22 weeks of gestation fetal tissues (*rows*). Higher transcription (Z-scores) are shown in green-yellow, lower expression and no expression is shown in purple and gray, respectively. Higher than average expression of I_γ4_ is observed in all tested tissues. Data from this figure was obtained from study SRP055513 [31].

### CSRnc transcription in cancer varies according to cancer type and is likely to depend on the degree of B cell tumor infiltration

The TCGA project data was used as a reference to study CSRnc transcription in a wide variety of human cancers. As for GTEx, CSRnc transcription (I_H_ average log_2_ RPKM distribution) was analyzed across 33 cancer types (Figure 8). I_H_ transcription in diffuse large B cell lymphoma (DLBCL) and acute myeloid leukemia (AML) were the highest. In general, CSRnc transcription in neoplastic tissue mimicked its non-neoplastic counterparts, being high in lung, stomach, and testicular germ cell carcinomas. Conversely, CSRnc transcription was low in glial cell and skin cancers (Figure 8).

**Figure 8.**
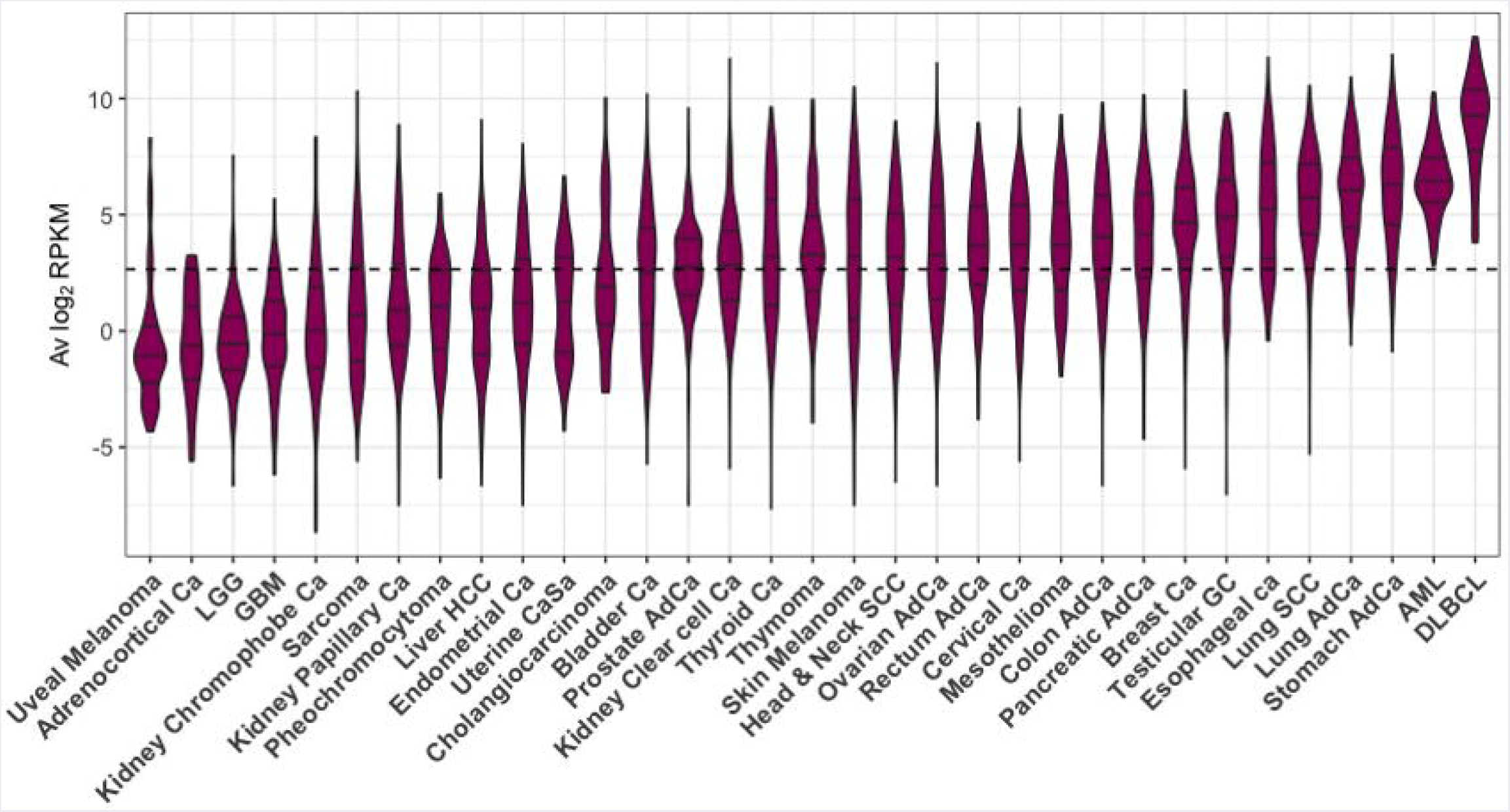
CSRnc transcription in cancer. RNA-seq data from the TCGA project was used to analyze CSRnc transcription in 33 cancer types. Violin plots of average log_2_RPKM, ordered from left to right according to increasing median. Violin area is scaled to each tumor sample count. A dotted black line marks the average log_2_RPKM = 2.65.

A direct comparison between CSRnc transcription in healthy (GTEx) and neoplastic tissue allowed the identification of three distinct patterns (Figure 9): 1) Tumors where average CSRnc transcription is lower than in healthy tissue, such as in DLBCL, prostate, thyroid, liver and colon cancer (Figure 9 A-E). 2) Tumors where average CSRnc transcription was higher than its healthy tissue counterpart, such as breast, rectum, testicular germ cell, pancreas carcinomas, ovarian cystadenoma and skin melanoma (Figure 9 G-L). 3) Tumors with no difference in CSRnc transcription between healthy and neoplastic counterpart, such as stomach and esophagus (**Additional file 1: Figure S10**). Interestingly, using healthy kidney cortex as reference for kidney tumors, CSRnc transcription varied according to cancer type, CSRnc transcription was significantly lower in chromophobe and papillary carcinomas, but not in clear cell carcinoma. (Figure 9F). In adrenal gland, a similar pattern was observed, with lower CSRnc transcription in adrenocortical carcinoma, but not pheochromocytoma (**Additional file 1: Figure S10**).

**Figure 9.**
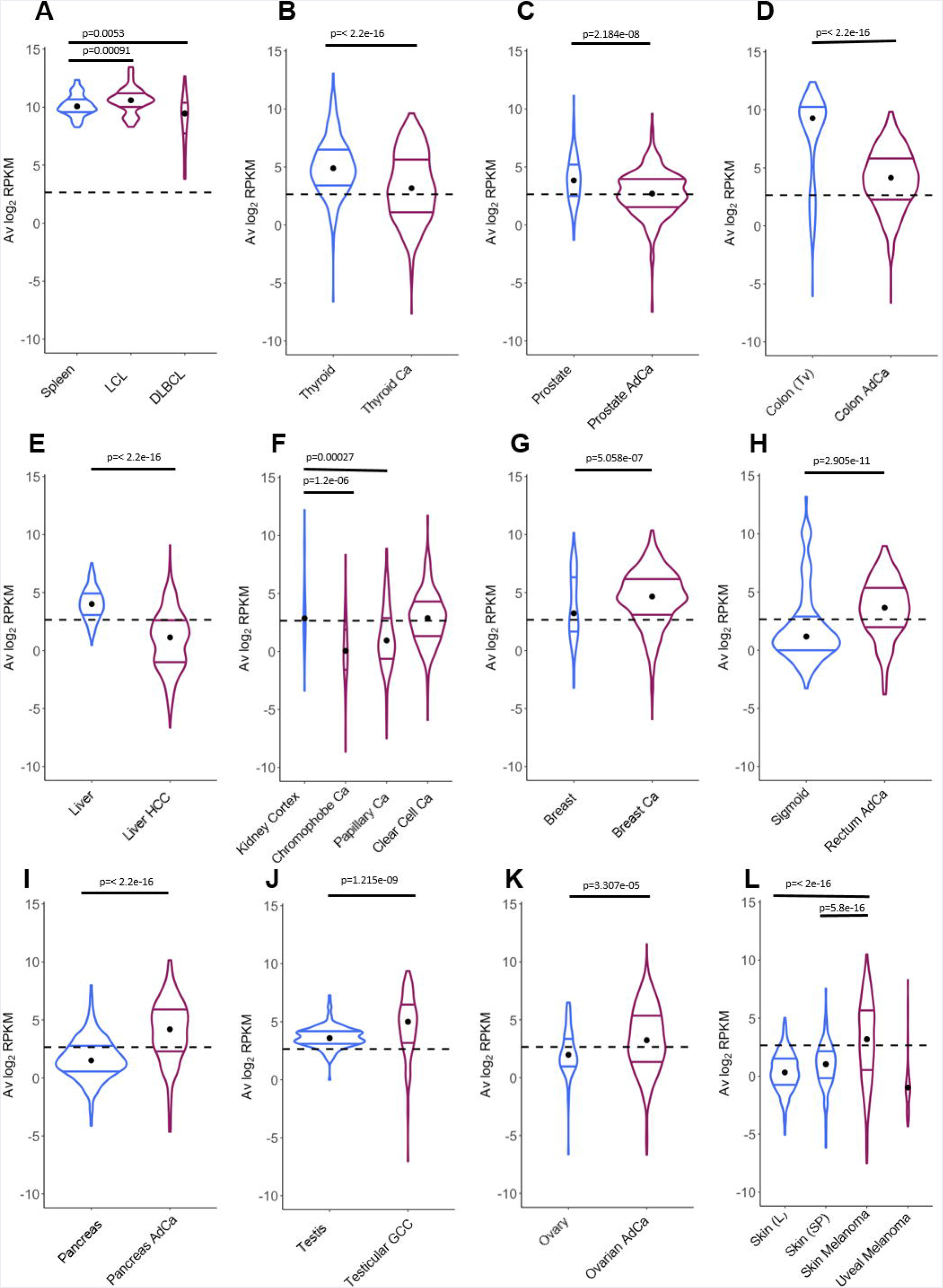
Comparison of CSRnc transcription in healthy tissue and its tumoral counterpart. Violin plots of the average log2RPKM distribution in healthy tissue (*blue*) in comparison with its cancer tissue counterpart (*purple*). Violin area are not scaled to sample count. Median (*black dot*) and quartiles are shown for each violin. Dashed black line marks the mean average log_2_RPKM (2.65). A reduction of CSRnc transcription in tumors was observed in **A-E**. An increase of CSRnc transcription in tumors vs. healthy tissue was observed in **G-K**. Some types of kidney cancers and melanomas showed an opposite patterns regarding the healthy tissue counterpart (**F** and **L**). The conducted statistical test were Wilcoxon rank sum test with continuity correction for two-sample comparisons, and Kruskal-Wallis test with *post hoc* Dunn’s test correction for multiple comparisons.

The correlation between CSRnc and C_H_ transcription (**Additional file 1: Figure S7**) suggests that as for healthy tissues, CSRnc transcription derives from tertiary lymphoid infiltrates resulting from tumor-associated inflammation and the corresponding mucosal associated lymphoid tissues [32]. To address this question, we analyzed CSRnc transcription in a wide variety of tumor-derived cell lines in SRA. The majority of the representative tumor derived cell lines tested (i.e., lung cancer A549 cells, breast cancer MCF-7 cells, and colon cancer HCT116 cells), were depleted in samples expressing CSRnc RNA (**Additional file 2: Table S2**), indicating that CSRnc transcription in cancer derives from infiltrating B cells and not the neoplastic cells *per se*. Nevertheless, using this approach, CSRnc transcription in cancer cells *in situ* cannot be ruled out.

### CSRnc transcription is altered in certain infectious conditions and a Ia-Ia2 transcriptional signature is associated with pediatric Crohn’s disease with deep ulceration

The SRA dataset represents the most diverse collection of data regarding methodological approaches and subjects of interest and represents a useful source of data related to diverse malignancies, as well as infectious and non-infectious inflammatory pathology. Using MetaSRA Disease Ontology annotations [33], we identified significant enrichment of acute AML, breast and lung cancers in SRA samples with CRSnc transcription, confirming our observations derived from TCGA data analysis.

Moreover, the SRA dataset allowed us to identify enriched CSRnc expression in infectious diseases such as diarrhea, brucellosis and malaria. However, in all cases, peripheral blood samples were used for these experiments. To distinguish if the enrichment was due to increased expression, rather than for the inherent enrichment observed in blood (Figures 4 and 5), we performed differential expression analysis comparing experimental groups provided by SRA metadata. Significant reduction of CSRnc transcription was detected in enterobacteria, but not in rotavirus diarrheal disease (SRP059039), malaria (SRP032775) [34], brucellosis and leishmaniasis (SRP059172) (**Additional file 2: Table S3**). SRA data from influenza vaccination (SRP020491) [15] was consistent with our own experimental data demonstrating no significant changes in CSRnc transcription 7 days postvaccination, regardless the plasmablast migration wave to peripheral blood at that time, indicating that CSRnc transcripts observed in peripheral blood do not derive from recently class-switched plasmablasts (Figures 2L-O). CSRnc transcription in response to influenza vaccination contrasted with the observation that natural infection with H7N9 [35] (SRP033696), in which we observed changes in coding and CSRnc transcription (**Additional file 2: Table S3**).

The SRA dataset also showed enrichment for autoimmune disease terms such as systemic lupus erythematous and autoinflammatory diseases like inflammatory bowel disease. No significant differential expression was observed in peripheral blood transcriptomes in systemic lupus erythematosus patients compared with healthy subjects (SRP062966) [36] (**Additional file 2: Table S3**). However, a study comparing the ileal transcriptome in pediatric Crohn’s disease (CD) with and without deep ulceration, and with or without ileal involvement (SRP042228) [37], revealed an increased significant expression for I_μ_, I_δ_ and I_α2_ in patients with deep ulcerated CD, regardless the presence of macroscopic or microscopic inflammation (**Figure 10 and Additional file 2: Table S3**).

**Figure 10.**
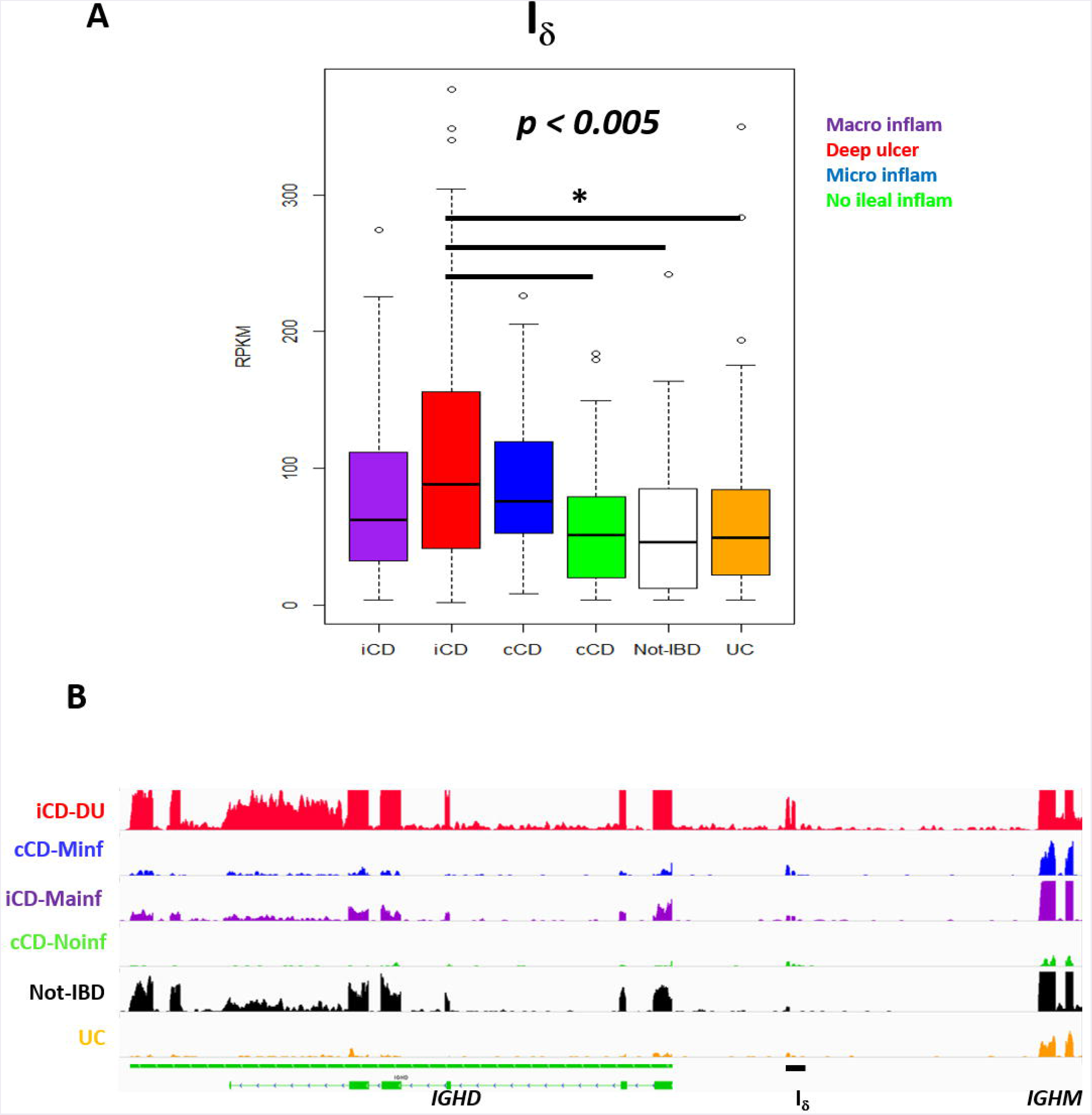
Increased Is transcription in ileal mucosa of Crohn’s disease. Pediatric Crohn’s disease patients from project SRP042228 were classified in two groups according to ileal mucosa involvement (iCD) on non-involvement (cCD) [37]. Patients were further classified according to the degree of ileal mucosa inflammation. Ileal mucosa from noninflammatory bowel disease (not-IBD) and ulcerative colitis (UC) were used as controls [37]. **A**) Boxplot showing increased Id transcription (RPKM, *y axis)* in deep ulcerated Crohn’s disease with ileal involvement (one factor ANOVA, Tukey multiple comparisons of means. P < 0.005). **B**) Transcriptional landscape of the *IGHD-IGHM* locus in human chromosome 14 showing increased I_δ_ transcription in representative samples of deep ulcerated iCD *(red track)*, cCD with microscopic inflammation *(blue track)*, iCD with macroscopic inflammation *(purple track)*, non-IBD *(black track)* and UC *(orange track)*.

## Discussion

We performed an integral, systematic analysis of human CSRnc transcription using an experimental approach and datamining of a large and diverse public RNA-seq dataset supported by a previously described resource, *recount2* [12]. The precise I-exon boundaries for every class were undefined and are not annotated in current human genome version (GRCh38.p12). Among hematopoietic-derived cells, CSRnc transcription was specific for B cells, and consistent with previous findings was present in naive as well as GC-B cells [5].

The present study has certain limitations. CSRnc transcription is required for CSR, however it does not prove that CSR is actively taking place. Among the most determinant factors influencing the amount of CSRnc transcription observed in an RNA-seq sample is the relative amount of B cells expressing a particular I-exon, as suggested by the high Z-score correlation between CSRnc and its corresponding C_H_ transcript (**Additional file 1: Figure S7**). We propose that the higher non-coding/coding Z-score ratio indicates a higher proportion of B cells undergoing a particular CSR event. Other factors influencing the amount of CSRnc transcription is that switch circles are transcriptionally active [38], and our current analysis cannot differentiate if CSRnc transcripts derive from chromosomal or switch circle transcription. Switch circles are non-replicating episomes that decay with B cell division [39-41]. Thus, B cells undergoing cell division at high rate (i.e., GC B cells) should dilute the amount of circle templates in a greater extent than non-dividing B cells.

A novel transcribed element (I_δ_) located in the *IGHM-IGHD* intergenic region was identified, which overlaps with a previously described repeat Σ_μ_ region, downstream of an atypical switch region (σδ) involved in non-classical CSR to IgD in mice [42] and humans [43-45]. The Σμ region was originally described as mediator of μ–δ CSR by homologous recombination in myeloma cell lines [22, 23]. The biological role of Σμ is uncertain, because IgD+ EBV-transformed cell lines and human tonsillar B cells undergo μ–δ CSR by non-homologous recombination using σδ as acceptor switch region [43, 45], in an AID-dependent fashion [44]. Here, we demonstrate that I_δ_ (Σμ) can be actively transcribed (Figure 1B). Nonetheless, I_δ_ as an acceptor I-exon does not fit into the general model of CSR, because it is located downstream of the σδ region. Further research is required to elucidate if I_δ_ transcription is involved in the non-classical μ–δ CSR.

An important motivation for this work was the identification of early transcriptional signatures in blood that correlated with the strength and quality of the humoral response, and in particular with the GC response. Interestingly, Hepatitis B and or Tetanus/induced I_μ_, and I_γ3_ transcription in peripheral blood at day 14 post-vaccination, but not when plasmablasts peaked (day 7 post-vaccination), suggesting that CSRnc transcription may not be the result of plasmablast mobilization to peripheral blood. Moreover, we observed increased I_γ1_ and I_α1_ transcription in natural H7N9 infection, but not upon influenza vaccination (**Figure 2, Additional file 2: Table S3**), or in rotavirus infection (**Additional file 2: Table S3**). The differential response observed upon influenza vaccination and natural rotavirus infection in contrast to HBV/tetanus-diphteria vaccination and H7N9 infection may be the result of the common repeated exposure to seasonal influenza or rotavirus, which would reactivate IgG+ memory B cells and in the absence of CSR [46].

Although I_H_ transcription is highly inducible upon activation (Figure 2A-D), GTEx and SRA mining revealed that I_μ_ transcription was higher than other I_H_. This is consistent with the current model of CSR, in which I_μ_ is constitutively transcribed under the control of the μ intronic enhancer (E_μ_), and its transcription is required for CSR regardless of the acceptor class [47]. E_μ_ participates in chromatin remodeling of the IgH locus during lineage commitment and VDJ recombination [48, 49], and its deletion impairs B cell development, I_μ_ transcription and CSR [50].

We identified distinctive CSRnc transcriptional patterns related to known immunological functions, such as I_μ_ and I_α2_ co-expression in MALT-rich organs, where active IgA secretion takes place. Of particular interest is cluster Z1, defined by an I_γ1_, I_γ2_ and I_γ4_ transcriptional signature. Z1 cluster was the only cluster with higher expression of I_γ4_, and was characteristic of the testis, thyroid gland and visceral adipose tissue (omentum), but not subcutaneous adipose tissue. Visceral adipose lymphoid B cells are immunologically active cells implicated in adipose tissue homeostasis that may play important pro-inflammatory roles associated with metabolic syndrome and obesity [51]. The expression of I_γ1_ and I_γ4_ transcriptional signature in testis and thyroid is unexpected, because they are regarded as immune-privileged sites devoid of secondary/tertiary lymphoid organs [52]. Similarly, higher I_ε_ transcription in the thyroid gland is a striking finding worthy of further research due to the common association of atopic disease with autoimmune thyroiditis [53]. At present, we do not know if CSR to IgG_4_ or IgE is taking place, however further research is required given that IgG_4_ is an atypical Ig that lacks Fc-mediated effector functions [54] and IgE could be implicated in autoimmunity. Furthermore, I_γ4_ transcription in different fetal tissues suggests that its transcription is not limited to lymphoid tissue, may be a common feature of multipotency and is not necessarily coupled with C_H_ transcription, suggesting additional functions beyond CSR.

A major goal of the GTEx project was to identify the role of genetic variation in gene expression as a quantitative trait. Tissue enrichment in two clusters with qualitatively different I_H_ transcriptional pattern such as in liver (Z3 and Z6), testis (Z1 and Z2), terminal ileum (Z5 and Z9) and whole blood (Z5, Z7 and Z9) indicate CSRnc transcriptional heterogeneity in tissue donors involved in the GTEx project. This could result from different tissue microenvironments (i.e., cytokine/chemokine milieu, microbiota and environmental stimuli) as well as genetics, which may modify CSRnc transcription patterns and possibly CSR itself. A recent study identified four SNP’s in the human IgH locus presumably involved in CSR that affect immunoglobulin levels [55]. Their implication in modifying CSRnc transcription warrants further investigation.

The study of CSRnc transcription in cancer is of particular interest for several reasons: 1) The anti-tumor response is largely mediated by the presentation of tumor neoantigens to T cells and Treg balance. However, antibody-mediated anti-tumor activity can be achieved by antibody - dependent cytotoxicity or other mechanisms [32]. 2) The presence of ectopic (or tertiary) lymphoid structures (ELS) [32, 56] and higher densities of infiltrating B cells and T follicular helper cells correlate with improved survival in lung [57], breast [58] and colorectal carcinoma [59]. 3) The tumor microenvironment, including certain cytokines may modify CSR patterns in infiltrating and ELS B cells, regardless the antibody effector function. 4) AID activity is a known contributor to off-target mutagenesis and genomic instability in B cell malignancies [60]. Aberrant AID and CSRnc transcription in non-lymphoid tumor cells could potentially contribute to cytidine deaminases - mediated kataegis [61].

We have found that average I_H_ expression in certain tumors analyzed in the TCGA project resembles their non-neoplastic counterpart, however some cancer types have significantly less CSRnc transcription, whereas others show the opposite. Most of the evidence we have gathered so far indicate that the origin of I_H_ transcription is in the tumor infiltrating and ELS B cells, rather than the tumor cells *per se* (**Additional file 2: Table S2**). Thus, differences in I_H_ transcription in cancer may be the result of immune editing [62], which may alter the amount and activation state of the infiltrating and ELS B cells. Based on gene expression signature clustering, all non-hematologic cancers of the TCGA project were classified into six immunologically distinct subtypes with distinctive somatic aberration patterns, tumor microenvironment including the amount and cell type infiltration, and clinical outcome [63, 64]. The relation between these six immunological subtypes with CSRnc transcription pattern may help to understand the elusive role of infiltrating B cells in the progression of different cancer types [32].

Despite the limitation of relying on public data when often the submitter researcher chooses to submit the minimal requirements of sample metadata, the SRA represents an enormous source of RNA-seq data from a highly diverse type of studies. In contrast to the standardized methodological criteria and metadata collection protocols used by the GTEx and TCGA consortiums, higher methodological variability in SRA data is expected, limiting inter-study comparisons. Nevertheless, we were able to identify a I_μ_- I_δ_- I_α2_ signature in ileal mucosa of Crohn’s disease in a treatment-naive pediatric cohort [37]. I_μ_ and I_α2_ is somehow expected as the result of predominant IgM to IgA CSR on mucosal tissue (Figures 4-6), and its exacerbation due to increased tissue B cell infiltration in response to inflammation [37]. However, the increased I_δ_ transcription, particularly in CD associated with deep ulceration is an intriguing finding (Figure 10). Serum IgD levels are elevated in patients with CD [65] and other autoinflammatory syndromes [44, 66]. A high proportion of μ–δ switched IgD+ cells bare autoreactive and poly-reactive specificities [67, 68], a feature shared with “natural autoantibodies”, which are reactive against bacterial wall components and may provide natural immunity against bacterial infection [69]. In human respiratory mucosa, μ–δ switched IgD^+^ B cells mediate the innate-adaptive immunity and inflammatory cross talk [44]. Mice incapable of undergoing classic CSR due loss of function of 53BP1 have an intestinal microbiota-dependent elevation of IgD serum titers and increased μ–δ switched IgD+ B cells [70]. Direct experimental testing is required to elucidate the role of I_δ_ transcription, its role in μ–δ CSR and its implications in healthy and inflamed mucosae.

## Conclusions

We have performed an unbiased analysis of the transcriptional landscape of the human *IGH* locus using a vast public RNA-seq dataset. Our observations agree with previous findings regarding constitutive CSRnc transcription in naïve B cells and its upregulation upon activation. We provide a detailed analysis of CSRnc transcription in healthy tissue. As expected, CSRnc transcription correlated with the amount of associated lymphoid tissue, however, novel transcriptional signatures involving I_γ4_ or I_ε_ were found in testis, pituitary, thyroid and visceral adipose tissue were identified. Changes in CSRnc transcription between healthy and tumor tissue were also found, likely as a result of immune editing. A novel transcribed element within the *IGHM-IGHD* intron termed I_δ_ was discovered and highly expressed in ileal mucosae of pediatric Crohn’ s disease patients. Overall, this study highlights the importance of open access data for discovery and generation of novel hypothesis amenable for direct testing, and is a great example of the potential of the *recount2* dataset to further our understanding of transcription, including regions outside the known transcriptome.

## Materials and methods

### Vaccination of human healthy volunteers

Pre-immune (day 0), day 7, 15, 30 and 180 post-vaccination peripheral blood samples (18 mL) were obtained by venipuncture in 2 x 8 mL Vacutainer^®^ CPT^™^ tubes from healthy volunteers vaccinated with Hepatitis B and/or Tetanus toxoid/Diphteria (n = 16), or Trivalent Influenza Vaccine during season 2013-2014 (A/California/7/2009 (H1N1) pdm09; A(H3N2) A/Victoria/361/2011; B/Massachusetts/ 2/2012) (n = 18). Written informed consent was obtained from each volunteer in each blood sample draw. All procedures in human subjects were performed after Institutional Review Board approval from the National Institute of Public Health (CI: 971/82-6684). Plasma and PBMCs were isolated according to the manufacturer’s instructions, aliquoted and stored at -80°C and liquid N2, respectively. Total RNA was extracted from PBMCs with TRIzol and stored at -80°C until used.

### Quantitation of plasmablasts by FACS

Cryopreserved PBMCs were thawed at 37° C and resuspended in RPMI 10% FBS, washed with PBS 1X and fixed with 1% paraformaldehyde for 20 min at room temperature. After washed with FACS solution (PBS 1%, sodium azide 0.05% and 2% FBS), cells were incubated for 30 min at 4 °C with the following antibody cocktail: anti-CD3 PerCP/Cy5.5 (clone SKY; Biolegend; 344808), anti-CD19 FITC (clone HIB19; Biolegend; 302206), anti-CD20 PE Cy7 (clone 2H7; Biolegend; 302312), anti-CD27 APC (clone O323; Biolegend;356410) and anti-CD38 PE (clone HIT2; Biolegend; 980302). Flow cytometry analysis was performed in a FACS Aria II (BD Biosciences, San Jose, CA, USA). Doublets and CD3+ events were gated out. Plasmablasts were defined as CD3^-^/CD19^+^/CD207CD27^+^/CD38^+^. 500-1000 plasmablasts were acquired per sample. Analysis was performed using Flowjo software (TreeStar).

### B cell *in vitro* stimulation

PBMCs were isolated by Ficoll-Paque^™^ density gradient from blood bank buffy coats. B cells were enriched through negative selection using B cell Isolation Kit II (MACS, Miltenyi). 1×10^6^ B cells were seeded per well on 6-well plate incubated in RPMI medium supplemented with 10% FBS, streptomycin and penicillin at 37°C with 5% CO_2_. Two activation conditions were stablished at different time points (3 and 6 days post-activation: Germinal center-like activation (GC-like), 1μg/ml anti-human CD40 (G28.5), 5μg/ml CpG ODN 2006 (Invivogen) and 25ng/ml recombinant IL-21 (eBiosciences). For T-independent activation, 5μg/ml CpG ODN 2006 (Invivogen), 0.05%*S.aureus* Cowan (Pansorbin; Calbiochem), 5ng/ml Pokeweed Mitogen (Sigma).

### qRT-PCR of CSRnc transcripts

Total RNA from PBMCs was extracted using TRIzol (Invitrogen). The integrity of the RNA was measured with Agilent RNA 6000 Nano. SuperScript^™^ III One-Step RT-PCR(Invitrogen) was used for reverse transcription and amplification. Quantitative PCR of CSRnc transcripts for *IGHM, IGHG1*, IGHG3 and AID gene was performed using specific primers and TaqMan probes(IDT). The primers and probes used to quantify the CSRnc transcripts are detailed in Table 3. Amplification of *HPRT* with PrimeTime^®^ Predesigned qPCR Assays was performed as the reference gene. The fold difference was calculated using 2ΔΔCT, using resting enriched B cells as calibrator and non-B cells as negative control. *HPRT* was used as normalizer for every condition.

**Table 3.**
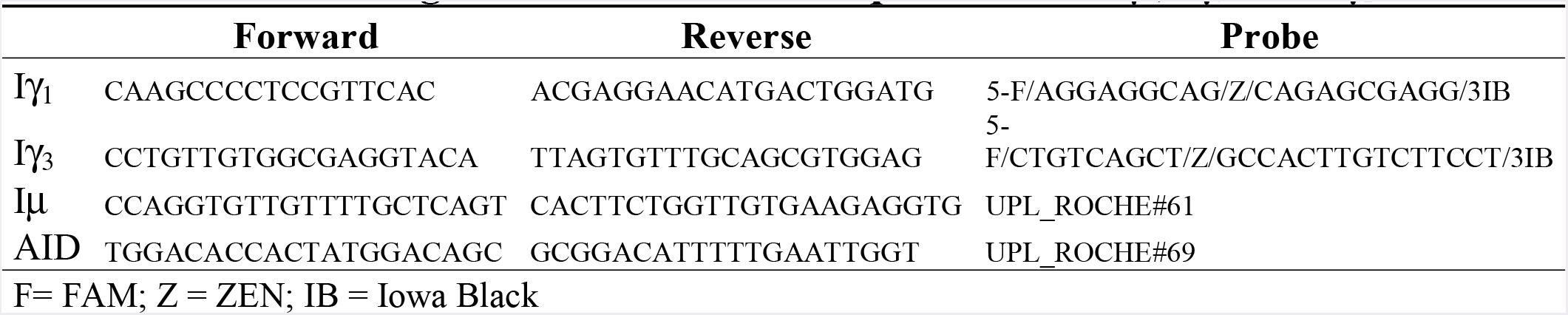
Oligonucleotides used for qRT-PCR of I_μ_, Iy_3_ and Iyi

### CSRnc transcription boundaries definition

Currently, CSRnc transcripts and switch regions (S) are not mapped as such in the current version of the human genome (GRCh38). I_H_ are upstream of the corresponding switch region (S_H_), thus we first mapped S regions based on the frequency distribution of the AGCT motif in 500 bp bins along the whole *IGH* locus (105,583,700 – 105,863,000) [3]. *recount2* is an online resource consisting of normalized RNA-seq gene and exon counts, as well as coverage *BigWig* files (https://jhubiostatistics.shinyapps.io/recount/) that can be programmatically accessed through the R programming language [71]. To map I_H_, metadata from all SRA projects contained in *recount2*, as well as through the SRA-Run selector engine were used to identify RNA-seq samples and samples performed using purified B cells. The corresponding *BigWig* files were downloaded using the *recount* Bioconductor package and mapped read counts were visually inspected with Integrative Genomics Viewer [72]. CSRnc regions were delimited according to an expression consensus from projects described in **Additional file 2: Table S1**.

### CSRnc and IGH transcription quantitation

*recount2* was used to extract read counts from each of the nine C_H_ constant region coding genes (*IGHM, IGHD, IGHG3, IGHG1, IGHA1, IGHG2, IGHG4, IGHE* and IGHA2), as well as from the corresponding CSRnc I exon coordinates as a *GRanges* object [73]. The log_2_-transformed C_H_ (coding) and CSRnc RPKM average per *sample* was used as an approximation of abundance of transcription. For CSRnc transcription, log_2_ RPKM per *sample* average adopted a quasi-normal distribution with a mean of 2.65 log_2_ RPKM (SD ± 3.79), which corresponds to 6.29 RPKM. As an initial exploration to which tissues and in which diseases CSRnc transcription takes place, we used the log2 RPKM average as a cut-off to define “high” expression (> 2.65 log_2_ RPKM) or “low” expression (< 2.65 log_2_ RPKM). The mean C_H_ transcription average log_2_ RPKM was 7.82 (SD± 5.16). Given the difference between coding and CSRnc transcription, and to address the relative expression between coding and CSRnc transcription for each Ig loci, coding and CSRnc log_2_RPKM values for each Ig gene were standardized by transformation to Z-scores.

### SRA RNA-seq samples Ontology mapping

Although all RNA-seq samples in TCGA and GTEx follow a homogeneous ontology categorization, metadata associated to SRA projects is widely heterogeneous and commonly insufficient. To obtain a more homogenous categorization of nearly half of our dataset, we used disease [33] annotations retrieved from MetaSRA, version 1-2 [74].

### Enrichment test

To define CSRnc transcription profile variation in healthy tissue (GTEx dataset), we performed tissue sample enrichment analysis according to Z-cluster using a two sided Fisher’s Exact Test. The H_0_ is that there is no difference in the probability distribution between Z clusters and tissue. A 2 x 2 contingency table was built for each tissue between the number of samples belonging to a given Z cluster and the remaining samples not belonging to that cluster. A two-sided Fisher’s test was performed with the R function fisher.test (c, alternative = “two.sided”). P value adjustment with the Benjamini-Hochberg method was performed to correct for multiple testing using the R function p.adjust(p, method = “bh”, n = length(p)). A False Discovery Rate (FDR) < 0.01 was considered as a significant enrichment.

### CSRnc transcription profile clustering

To define CSRnc transcription profiles, Z-scores for each I-exon were subjected to *k*-means clustering using the Cluster software 3.0 [75]. Ten clusters were generated, with 100 iterative runs and Euclidean distance as distance metric. Clustered data was visualized with java Treeview 3.0 (http s:// sourceforge.net/projects/ jtreeview/).

### Differential expression analysis

Differential expression of coding IGH and CSRnc transcripts was analyzed using the functions lmFit() and eBayes() from the *limma* R package v3.34 [76]. If technical replicates were present for a given study, the induced correlation was adjusted for using the duplicationCorrelation() function from *limma*. The resulting Bonferroni-adjusted p-values less than 0.05 were determined to be statistically significant. We used Bonferroni instead of FDR given that we tested 10 regions instead of the usual number of thousands of genes.

## Additional file 1

**Figure S1**, Distribution of the number of samples per RNA-seq project analyzed. **Figure S2**, CSRnc transcription is B cells-specific. **Figure S3**, CSRnc transcription pattern in whole blood is similar to peripheral blood sorted CD19^+^ B cells. **Figure S4**, Epigenetic marks in I_δ_. **Figure S5**, CSRnc transcription according to I_H_ and project dataset. **Figure S6**, CSRnc transcriptional profiles identified by *k*-means clustering of the recount2 dataset **Figure S7**, Correlation between CSRnc I_H_ transcription and coding C_H_ transcription. **Figure S8**, CSRnc and C_H_ coding transcription in Bone marrow lymphoid precursors. **Figure S9**, CSRnc and C_H_ coding transcription in thymic lymphoid precursors. **Figure S10**, Comparison of CSRnc transcription in healthy tissue and its tumor counterpart (PDF).

## Additional file 2

**Table S1**. Selected SRA projects used to map I_H_ boundaries. **Table S2**, CSRnc transcription analysis in cancer cell lines. **Table S3**, Differential expression analysis in I_H_ and C_H_.(XLSX).

## Ethics approval and consent to participate

The participants in this study did so voluntarily after written consent. The study was approved by the INSP Institutional Review Board (CI: 971/82-6684).

## Acknowledgements

We would like to thank Analí Migueles and Everardo Millan for their support in data preprocessing, and Fernando Riveros Mackay for differential expression analysis; Leopoldo Santos-Argumedo for kindly donating αCD40 hybridoma G28.5), Martha Patricia Rojo for support on flow cytometry; Tomás Salmerón Enciso, Yvonne Rosenstein, Fernando Esquivel and José Moreno for academic input, and Menno Van Zelm for critically reviewing the manuscript. HKM is a PhD student of Programa de Doctorado en Ciencias Biomédicas, UNAM, and was supported by a CONACyT scholarship.

## Abbreviations

AID: Activated-Induced Cytidine Deaminase
CD: Crohn’s disease
CSRnc: Class Switch Recombination non-coding
CSR: Class Switch Recombination
CG: Germinal Center
C_H_: Heavy chain Constant
ELS: Ectopic Lymphoid Structures
EBV: Epstein Barr Virus
GTEx: Genotype-Tissue Expression Project
I_H_: I exon
MALT: Mucosal Associated lymphoid Tissue
PAMP: Pathogen-Associated Molecular Pattern
PB: Peripheral blood
SRA: Sequence Read Archive
RPKM: Reads Per Kilobase (transcript) Per Million (reads)
TCGA: The Cancer Genome Atlas.

## Authors’ contributions

This study was conceived and designed by HKM and JMB. The experimental procedures were performed by HKM, MOM, HVT and JMTS. Flow cytometry acquisition and analysis was conducted by HKM and LBA. RNA-seq data bioinformatics and statistical analysis were done by HKM, LCT and JMB, and supervised by AJ. HKM and JMB drafted the manuscript. LBA, LCT and AJ critically reviewed the manuscript. All authors read and approved the final manuscript.

